# Insecticide-Releasing LLDPE Films as Greenhouse Cover Materials

**DOI:** 10.1101/381426

**Authors:** Senem Avaz Seven, Ömer Faruk Tastan, Cüneyt Erdinç Taş, Hayriye Ünal, İkbal Agah Ince, Yusuf Ziya Menceloglu

## Abstract

The use of chemical pesticides is limited by several public health concerns regarding their toxicity levels and indiscriminate use. Nevertheless, they are still vital components of agricultural industry since no other competitive equivalents to chemical pesticides still exist in terms of efficiency. This study describes the preparation and biological assessment of an insecticide releasing plastic film for agricultural covering purposes. The formulation was prepared by incorporation of deltamethrin loaded, nano-sized halloysite nanotubes into polymeric films. Thermal, morphological, and mechanical properties of films were characterized by Differential Scanning Calorimetry (DSC), Scanning Electron Microscopy (SEM) and Universal Testing Machine UTM. Sustained release profiles of the films were evaluated by Thermogravimetric Analysis (TGA). Results reveal that deltamethrin was successfully loaded into halloysite nanotubes and nanotube incorporation enhances the elastic modulus of linear-low density polyethylene (LLDPE) films. In addition, films exhibit controlled release function of the active agent for 32 days. Bioassays of the nanocomposite films with varying deltamethrin doses tested on grasshoppers showed that the *LD_50_* values of the films are 1.85*x*10^−5^ g/cm^2^. Insecticidal activities of films were tested in greenhouse on *Medicago Sativa* plants contaminated with thrips and aphid. Nanocomposites are observed to repel mature aphids and kill young aphids and thrips.

## 1. Introduction

Insects, herbs, and fungus have been addressed as the most important hazards in agricultural sector. Since these species may cause up to 30-40% loss in product loss per annum, crop protection chemicals have the largest market share between agricultural products and is reported to be growing in long term and will dominate the competition with low cost and sustainable alternatives [1]. On the other side of the story, only 0.1 % of the pesticides applied in an agricultural area reach the target, while the excess amount spreads into the soil and damage the environmental health [2]. The term pesticide covers insecticides, fungicides, herbicides acaricides, and rodenticide or mollusicides. Excessive use of pesticides results in loss of biodiversity and decrease in invaluable species [3], water and soil pollution [4], development of pesticide resistance in plants [5], and numerous other public health risks.

Among the crop protection pesticides, insecticides dominate the pesticide products market by 44% [6] and is predicted to reach $16.7 billion, globally, by 2020 [7]. The field of pesticides application is limited by several public health concerns regarding the use of pesticides. One of the major concerns includes the indiscriminate use of pesticides. It is reported that at least 90% of the chemicals used is lost in soil and could not reach the area where pest control is crucial [8–10]. Indiscriminate use of insecticides in conventional agriculture poses important public and environmental health risks considering the hazardous effects of synthetic pesticides. For instance, the excess amount of pesticides due to indiscriminate use gives rise to pathogen resistance and decreases the biodiversity of soil, reduces nitrogen binding, and increases bioaccumulation of pesticides [11]. Nevertheless, use of pesticides is inevitable to increase the agricultural reproductivity.

Several plant-based botanical insecticides have been proposed so far, but these products could not gained significant commercial achievements [12]. On the contrary to the emerging technologies in the field of biopesticides [13], chemical crop protection agents are still the most cost-effective alternative in pest control today. Synthetic pesticides cause acute or chronicle health issues in consumers by accumulating on agricultural products. Thus, the accumulation of synthetic pesticides has become a subject of public health. Therefore, the search for novel chemical insecticides, or development of controlled release formulations of commercial insecticides as safer alternatives is ever increasing. Considering the fact that developing novel chemical pesticide takes about 8 years to the market [14], alternative formulations of the existing commercial insecticides, such as controlled release, may provide shorter term solution to this problem. As a synthetic insecticide, deltamethrin leads the insecticide market due to its effect broad range of insects. Deltamethrin is an important synthetic insecticide in terms of both its high efficiency and effective protection in very low concentrations. Mostly found in spray formulation in the market, deltamethrin is a relatively safe insecticide with an LD50 value of 431 mg [15]. In addition, deltamethrin is a stable material due to its high decomposition temperature (400 °C), therefore is suitable for melt compounding and blending purposes [16, 17]. Although it is strictly dependent on targeted pest type, World Health Organization (WHO) approved rate of use of deltamethrin suitable for cold or thermal fogging for mosquito control is 0.5-1.0 g/ha [18].

Integration of nanotechnology in pesticide delivery systems gave rise to various possibilities to develop environmentally friendly insecticide formulations. Application of nanotechnology on pesticide delivery systems is a relatively new subject and in the early stages of development. Nevertheless, it gained enough acceleration in the past decade to create a reformation in agricultural methods [19]. Potential applications of this technology include pest control, controlled pesticide release, agrochemicals and pathogen detection [20–22]. In the area of agricultural product delivery, nanoparticles are novel in terms of product efficiency increment. The current research trends reveal that with “smart” delivery systems, it is possible to develop controlled release formulations of agrochemicals similar to nano-drug applications [23]. The literature of controlled insecticide release and/or delivery mostly relies on other pesticide species, and to our knowledge, studies subjecting controlled insecticide release are not frequently found in the literature. There are several studies in the literature that focus on controlled release of deltamethrin. One of these studies involves a polymer-based system that releases deltamethrin where, a veterinary purpose deltamethrin-containing insecticide mixture was completely released from polymeric substance [24]. Another study on controlled release formulations of insecticides reports that biological char/biosilica based pesticide formulations exhibit high absorption capacity and stability [25]. The reason to this fact was explained with porous nature of the carrier nanomaterial.

In the area of plant protection products, the main role of nanomaterials is to increase the efficiency of active material that is absorbed by the plant [26]. An example to this is [27] where nanomagnetic materials were added to insecticide formulations to increase their bioactivity. On the other hand, nanomaterials are also employed to enhance the stability of the plant protection product. A recent study demonstrates that UV-stability of pesticide species is increased by addition of triazole-containing polyelectrolyte formulations [28].

As a natural bioavailable nanomaterial, halloysite has gained much attention mostly due to its tubular structure. Along with its nano-sized radius and contrast chemical and electrical properties of inner and outer lumens, this tubular structure makes halloysite nanotubes (HNT) a promising candidate in new industrial research areas. Potential applications of HNTs include microfiber filler materials [29], chemical carriers [30], and drug delivery systems by controlled or delayed release of the active material [31], and anticorrosive coatings [32]. Moreover, HNTs is found to be suitable to biomedical materials production thanks to their low toxicity [33]. In addition to all these, it is important to note that HNTs are also very convenient for nanocomposite applications due to its cost-efficient mass production.

A notable aspect of HNT is the inner lumen radius allows loading of various materials, including macromolecules, into the nanotubes [34–37]. In an early study [38], drug molecules were loaded into HNT by saturated antibiotic solutions or melt treatment, and controlled release properties were investigated. Here, loading was performed in a vacuum medium by exchanging entrapped air pockets in the inner cavities with drug molecules. Removal of air pockets was observed in the form of bubbles leaving the solution. Although the 15 nm thick wettable inner lumen of halloysite creates very high capillary effect, the vacuum medium was reported to be crucial for high loading efficiency.

HNTs usually follow Al_2_Si_2_O_5_(OH)_4._nH_2_O (n=4) stoichiometry [39]. Tube perimeter is composed of monoclinic crystal multilayers. The inner lumen can be filled by functional materials up to 20-30% of the tube’s volume by chemical etching [37]. In addition to controlled release function, the strength of HNTs can be enhanced 3-5% by addition of polymeric materials [40]. The outermost lumen of HNT is composed of silica with an electrical potential of around −30 mV (pH 4.8) [41]. While this potential is lower compared to pure silica particles (−50 mV), the difference can be explained by positively charged alumina inner wall. The outer charge enables HNT to remain stable in water for 2-3 hours. Loading anionic species into the nanotubes makes positively charged inner wall more neutral, therefore, allows obtaining more stable aqueous HNT solutions [41, 42]. This way, duration of controlled release is reported to take place up to 4-12 hours [43].

The reasons mentioned above makes HNT a very good candidate as a controlled release agent. Several active materials were incorporated into HNTS such as silver [44], pesticides [45] to perform antibacterial coating agents. Yet, new formulations releasing the active agrochemical should be developed for agricultural applications. Among the commercial greenhouse-covering materials, plastics are preferred due to their lower cost, lower weight and availability to mass production compared to glass alternatives. Commercially purposed greenhouse materials are usually composed of linear, low-density polyethylene (LLDPE) based films. Research trend in development of LLDPE based films as greenhouse cover materials is particularly towards addition of reflective materials into LLDPE films to improve their optical properties. For instance, infrared reflective material incorporated LLDPE films was prepared by melt blending and blown film extrusion methods [46]. Considering the fact that greenhouse-covering materials are subject to varying climate conditions, the prominent features expected from a candidate can be listed as mechanical strength, convenient optical properties and thermal stability.

Literature examples of additive incorporated polyethylene films focus on antimicrobial film applications. For instance, LLDPE films were incorporated with essential oils to produce controlled release antimicrobial films for food preserving applications [47, 48]. In a very recent study, it was demonstrated that the addition of halloysite nanotubes into high density polyethylene (HDPE) films promote crystal growth and enhance thermal and mechanical properties up to a certain halloysite concentration [49]. In another study, LLDPE films were incorporated with HNTs to introduce ethylene scavenging and barrier properties into films that promote food packaging applications [50].

This study aims to develop greenhouse cover materials from LLDPE-based films with controlled insecticide release function. For this purpose, commercial insecticides were loaded into naturally occurring HNTs, and then the controlled release material was incorporated into LLDPE film. Results suggested that the prepared film shows similar thermal and mechanical properties to neat LLDPE. Controlled release of insecticide was successfully demonstrated and the duration was recorded as 32 days.

## 2. Materials and Methods

### 2.1. Deltamethrin Loading into HNT

Halloysite nanotubes in 65 μm of agglomerate size were kindly provided by ESAN Eczacibaşi as purified and pretreated via ball mill homogenization. Granules of linear-low density polyethylene (LLDPE) were supplied from PETKİM Petrokimya A.Ş., with a melt flow rate of 2-3.5 g/10min (190 °C, ASTM D-1238), and a density of 0.918 - 0.922 g/cm^3^.

To prepare deltamethrin-loaded halloysite nanotubes (DMHNT), HNTs were dispersed in water (20%) and sonicated for 30 min. and excess amount of deltamethrin (DM) was added into HNT dispersion. The mixture was shear-mixed at 7500 rpm for 10 minutes. Once a stable dispersion was obtained, HNT-DM mixture was subjected to 1 mBar of pressure to remove the air bubbles inside HNTs. DM was loaded inside HNTs by bringing this mixture to atmospheric pressure. To ensure an efficient loading, this procedure was repeated three times. DM loaded HNTs were then washed and separated by ultracentrifugation (7000 rpm, 10 min.) and dried by lyophilization.

### 2.2. LLDPE/DM-HNT Film preparation

DMHNT loaded LLDPE films were prepared at Sabancı University Integrated Manufacturing Technologies Research and Application Center (SU-IMC). To prepare the DMHNT loaded LLDPE films (LLDPE/DMHNT) at different ratios; DMHNTs were dispersed in linear low-density polyethylene (LLDPE) films. For this purpose, HNTs were compounded with LDPE granules by thermomixing. The masterbatches were prepared by a laboratory scale high-shear thermo-kinetic mixer (Gelimat G1, Draiswerke, USA). The resulting compounds were blow moulded to thick films using a single-screw extruder (Ultra Micro Film Blowing Line, LUMF-150 with LE8-30/C, LabTech Engineering). DMHNT added PE films prepared in different concentrations were described in Table 1.

**Table 1.** Deltamethrin-loaded HNT contents of LLDPE films prepared.

| Code | DMHNT Content in LLDPE (%) | Dose (g/cm<sup>2</sup>) |
| --- | --- | --- |
| LLDPE/DMHNT-0.0 | 0.0 | 0 |
| LLDPE/DMHNT-0.1 | 0.1 | $1.12 \times 10^{-6}$ |
| LLDPE/DMHNT-1.0 | 1.0 | $1.12 \times 10^{-5}$ |
| LLDPE/DMHNT-3.0 | 3.0 | $3.37 \times 10^{-5}$ |
| LLDPE/DMHNT-5.0 | 5.0 | $5.16 \times 10^{-5}$ |
| LLDPE/DMHNT-10 | 10 | $1.12 \times 10^{-4}$ |

### 2.3. Thermal Characterization

Confirmation and quantification of active material loaded into HNTs and sustained release properties were evaluated by thermogravimetric analysis instrument (TGA, DTG60H, Schimadzu). Samples were subjected to a heating curve ranging between 30-1000 °C with a heating rate of 10 °C/min under nitrogen atmosphere. Results were evaluated using TGA collection software (TA60-WS).

Thermal properties of DM-HNT loaded LLDPE thick films were evaluated by Differential Scanning Calorimetry (DSC; TA2000, TA Instruments). For these measurements, samples were heated in a heating cooling cycle ranging between 25-200 °C, with a heating/cooling rate of 10 °C/min. Before the measurement, samples were subjected to a pre-heat treatment to eliminate any thermal history. DSC thermograms were then recorded to gather thermal parameters such as crystallization temperature (*T_c_*), melting temperature (*T_m_*), enthalpy of crystallization (*ΔH_c_*) and enthalpy of fusion (*ΔH_m_*) from the onset temperature, peak temperature of the DSC peak and area under the curve, respectively. Relative crystallinity (*X_c_*) values for varying DM-HNT content were then calculated by thermograms using the equation,

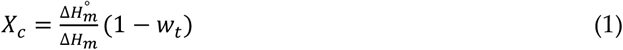

Where *ΔH°_m_* stands for the theoretical specific melting heat of 100% crystalline sample, and *w_t_* represents weight fraction of added DM-HNTs. For LLDPE samples, *ΔH°_m_* value was taken as 293 J/g [51].

### 2.4. Mechanical Performance Assessment

Mechanical properties of the prepared films were evaluated by Universal testing machine (UTM). For this purpose, dog-bone type specimens (*wxl=15×38 mm, grip distance=22mm*) of the thin films were prepared according to ASTM D-1708 standard. Tests were performed on a Zwick tester (Z100) under room temperature and humidity conditions. The crosshead speed was set as 12.5 mm/min (at L_0_=24 mm). At least three measurements were carried out per sample.

### 2.5. Evaluation of Morphological Traits

Morphological traits of LLDPE/DMHNT nanocomposites were evaluated by Scanning Electron Microscopy (SEM). To do this, sample surfaces were first sputter-coated with a thin layer of Pt/Pd and images were acquired by secondary electron and inLens detectors and a various gun voltage values ranging between 2-5 kV.

### 2.6. Dose-Response Bioassay

20 adult grasshoppers (*Locusta Migratoria*) (Mira company Antalya/Turkey) were used per dose and control with triplicates using independent insect cages. DM-HNT incorporated LLDPE films were adjusted to *10×10* cm per cage. Films were placed to the lid of the cages. Grasshoppers were fed with artificial diet. Dead insect numbers were counted every 24 hours until the end of the assay. Cages were maintained at *25 ±1* °C, and *16:8* light:dark photoperiod. The effects were evaluated up to 7 days in that high dose reached 100% mortality. The average of the three measurements was calculated and data were corrected using Abbott’s correction to calculate natural mortality [52]. *LC50* values (concentration causing %50 larval mortality) were calculated by log-probit regression using *SPSS 10.0* for Windows/Microsoft Excel program [53].

### 2.6. Insecticidal Activity

Once the DMHNT loaded LLDPE films were prepared, insecticidal activity was evaluated qualitatively at a Venlo type greenhouse built in Sabancı University, Istanbul [54]. The temperature was kept constant (*25* ± *2* °C in the daytime and at *18* ± *2* °C at night). For this purpose, *25×25* cm square samples were placed around *Medicago sativa* (alfalfa) plants that were contaminated with thrips and aphid. The samples were placed around *1 m^2^* of plant’s periphery.

## 3. Results

### 3.1. Confirmation of Deltamethrin Loading

DM loading was confirmed by thermogravimetric Analysis (TGA) and results were demonstrated in Figure 1. Neat HNTs and DM-loaded HNTs were subjected to a heating curve ranging between 30-1000 °C with a heating rate of 10 °C/min. DM loading was confirmed by the difference in total weight loss between unloaded and DM-loaded HNTs. Nanotubes mainly exhibit a two-step weight-loss. In the first step (>100 °C) HNTs lose their absorbed water content, while in the second step (450-550 °C) dehydroxylation of inner alumina sheet takes place. On the other hand, deltamethrin exhibit a weight loss in one abrupt step between 250-320 °C (data not shown here). Therefore, the weight loss ‘till 300 °C observed in DM-HNTs TGA curve arises from deltamethrin loaded inside HNTs. The difference in overall weight loss between unloaded and DM-loaded HNTs calculated as 17.9%, which corresponds to the amount of DM loaded into HNTs.

**Figure 1.**
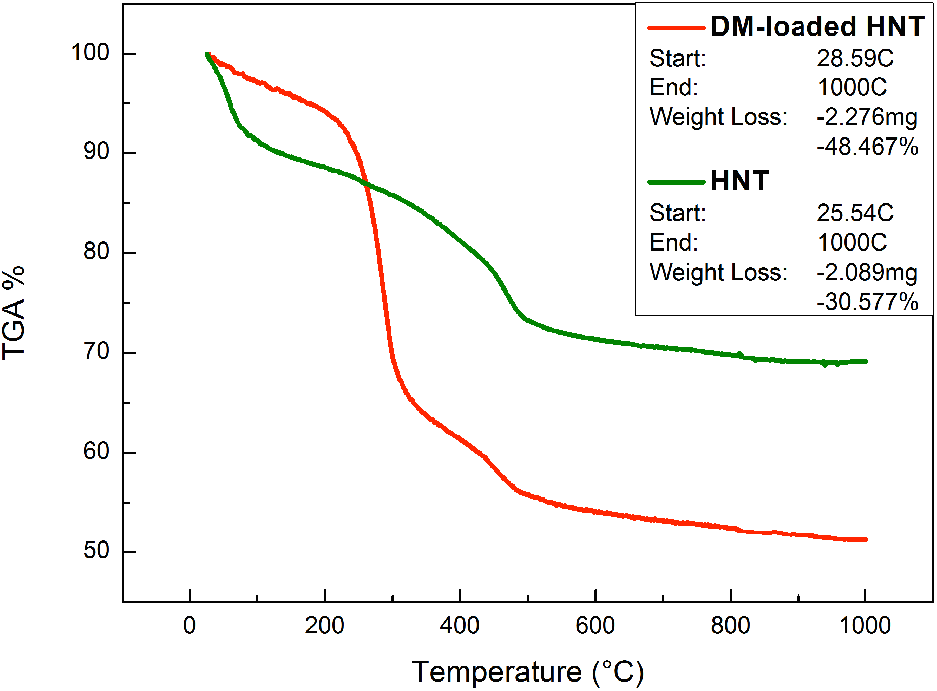
Thermogravimetric analysis of unloaded (green) and DM-loaded (red) HNTs.

### 3.2. Sustained Release Profile of Deltamethrin-Loaded Halloysite Nanotubes

Sustained release profile of DM loaded HNTs were investigated by termogravimetry. For this purpose, 5.22 mg of DM-loaded HNTs were kept at constant temperature (25 °C) and weight loss was monitored for 7 days. At the end of 7 days, the total weight loss is calculated as 3.98%. Isothermal TGA analysis of the sustained release profile is demonstrated in Figure 2.

**Figure 2.**
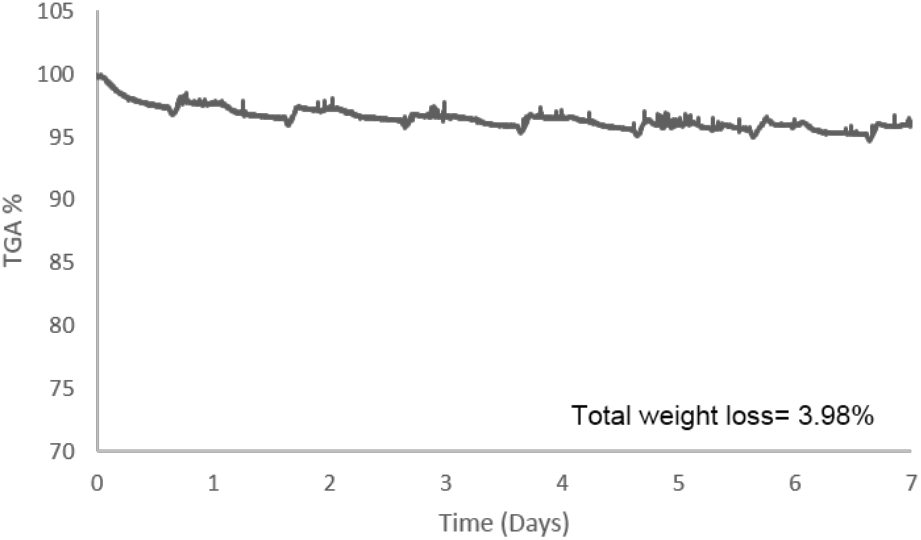
Isothermal thermogravimetric curve of DM-loaded HNT. Temperature was kept at 25 °C for 7 days.

In conventional agriculture practices, insecticide application is usually irregular. In a study [55], it was stated that the most frequently used conventional insecticides are applied 10-20 times annually. This means that a controlled release formulation should be actively releasing the agrochemical for 18-36 days. Recall that the total DM loading amount in this study was found as 17.9%. Assuming that HNTs are able to release all their content, DM-loaded HNTs sustain the release of deltamethrin for 32 days. Considering the finding that DM-loaded HNTs (10%) release 22.3% of deltamethrin per week, DM-loaded HNTs can be used as an insecticide agent for 32 days.

### 3.3. Thermal Properties

Thermal properties of neat and DM-HNT loaded LLDPE samples were also studied with DSC were (Table 2). From the data it is observed that melting and crystallization temperatures of neat LLDPE did not alter significantly upon DM-HNT loading. On the other hand, addition of 0.1-5% of DM-HNT into LLDPE films increase the crystallization enthalpy and percent crystallinity of LLDPE. This trend can be explained by the nucleating effect of polymer chains in the presence of nanoparticles [56, 57]. However, for 10% DM-HNT loaded LLDPE these values are observed to decrease in contrast with smaller concentrations. The decrease is attributed to the presence of aggregates of DM-HNT nanoparticles within LLDPE matrix.

**Table 2.** Thermal parameters of neat and DM-HNT loaded LLDPE.

| Sample | $T_m$<br>(°C) | $T_c$<br>(°C) | $\Delta H_m$<br>(J/g) | $\Delta H_c$<br>(J/g) | $X_c$<br>(%) |
| --- | --- | --- | --- | --- | --- |
| LLDPE/DMHNT-0.0 | 100.05 | 97.93 | 64.18 | 57.22 | 19.53 |
| LLDPE/DMHNT-0.1 | 100.31 | 97.85 | 62.64 | 59.41 | 20.07 |
| LLDPE/DMHNT-1.0 | 99.87 | 97.50 | 57.56 | 57.08 | 19.29 |
| LLDPE/DMHNT-3.0 | 100.23 | 98.16 | 67.33 | 63.13 | 20.90 |
| LLDPE/DMHNT-5.0 | 100.56 | 98.35 | 58.48 | 60.62 | 19.66 |
| LLDPE/DMHNT-10 | 100.84 | 98.35 | 57.73 | 56.58 | 17.37 |

### 3.4. Film Properties

DMHNT added PE films prepared in different concentration were subjected to tensile tests. Elastic modulus (E_T_), ultimate tensile strength (σ_max_), percent elongation at break (ε), and toughness (W) values of LLDPE/DMHNT films are demonstrated in Figure 3.

**Figure 3.**
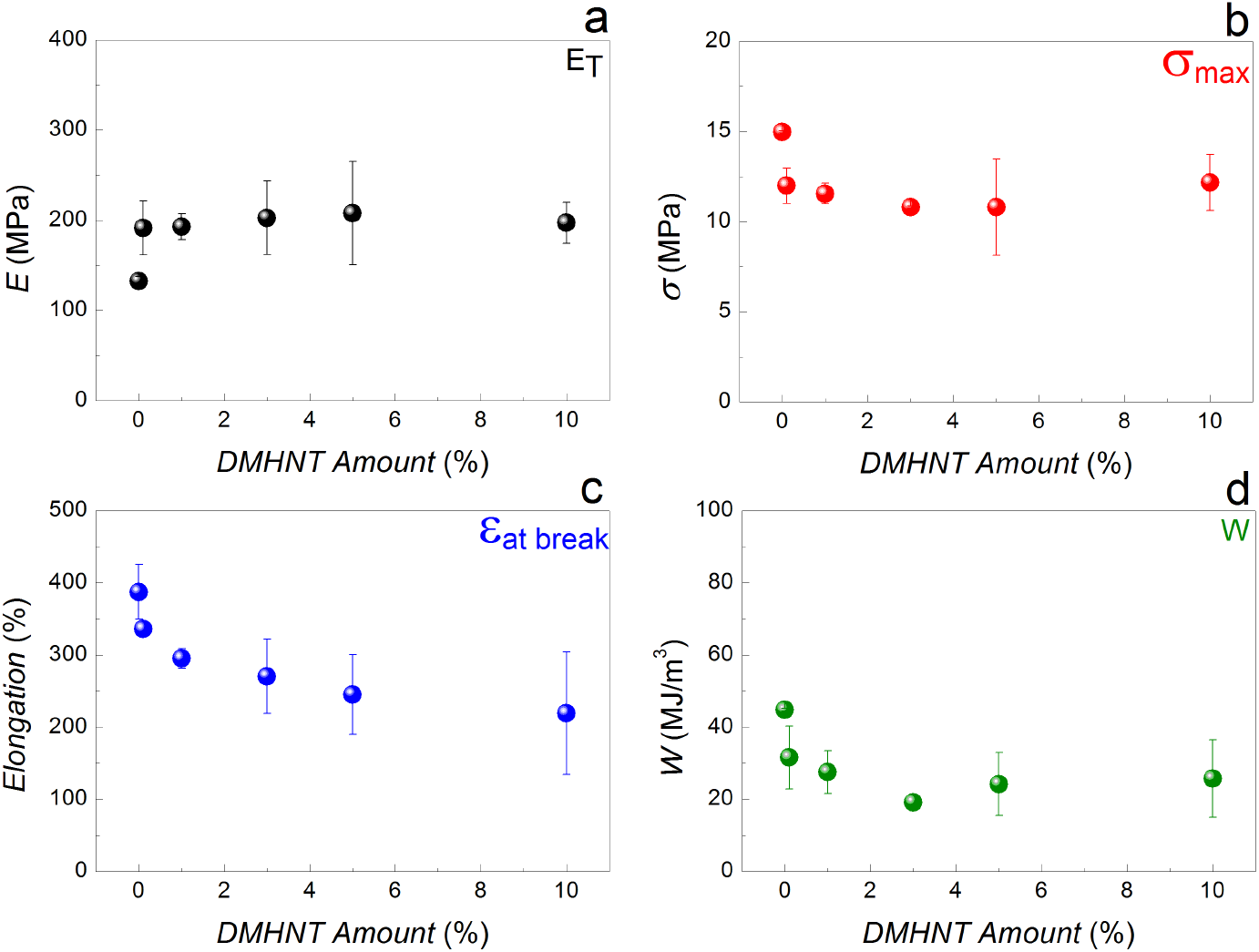
Tensile Properties of LLDPE/DM-HNT samples; elastic modulus (a), ultimate tensile strength (b), % elongation at break (c), and fracture toughness (d).

Mechanical tests reveal that DM-HNT incorporation enhances the stiffness of the films by an average of 40% (Fig 3-a), with a slight drop in ultimate tensile strength, ductility and toughness (Fig 3-b, c, d, respectively). The drop in tensile strength is attributed to the finding [58] that the agglomerates observed within the polymeric matrix due to HNT incorporation upon extrusion (Figure 4-d) act as stress concentrators. Although it is easy to address the extrusion process as the source of agglomeration, one should also note that the slipping of reinforced HNTs during mechanical test might also be the reason of agglomeration of HNTs and therefore the drop in tensile strength and other mechanical properties.

**Figure 4.**
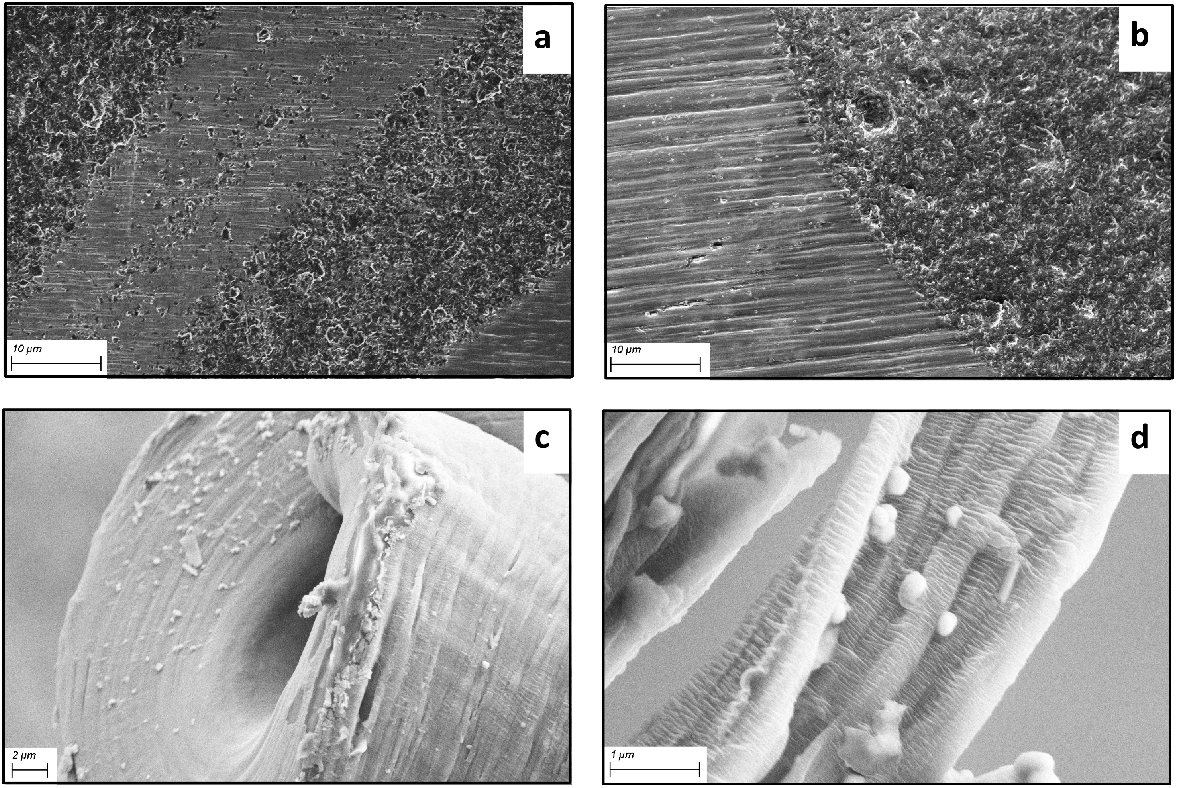
SEM images of neat LLDPE and its nanocomposites at several magnifications. InLens detector images of neat LLDPE (a), and 5%-DMHNT composite (b) were gathered at 2 and 3 kV, respectively; SEI detector images of 5%-DMHNT nanocomposite at different magnifications (c, d) gathered at 5 kV.

### 3.5. Morphology

Figure 4 shows morphological traits of neat and LLDPE/DMHNT nanocomposites obtained by SEM. Images were recorded from plastic deformation surface of neat LLDPE and 5% DMHNT (Fig 4-a, b; respectively), and fracture surface of 5% DM-HNT (Fig 4-c) tensile test specimen. In Figure 4 a and b, linear lines represent elastically deformed regions of neat and 5%-DMHNT LLDPE films, while plastic shear deformation is observed as rough surfaces. HNTs on the other hand, are observed as white dots (Fig 4-b). Although introduction of rigid nanoparticles is expected to deflect crack propagation and thus giving rise to increased strength, in this case the nanoparticle agglomerates promote premature tensile failure prior to crack [59]. Mostly uniform distribution of HNTs within 5% DMHNT is observed in Figure 4-b, c, and -d along with some small agglomerates.

### 3.6. Bioassay

DM-HNT loaded LLDPE films demonstrate promising activities against all the tested grasshopper species. The regression parameters of probit analysis and *LD50* for mortality of the larvae are presented in Figure 5. Data confirms that lethal 50 concentrations of films are 1.85*x*10^−5^ g/cm^2^, which lies between 1% and 3% doses applied. Table 3 shows the overall death rate results of 0.1, 1, 3, and 5% DM-LLDPEs, along with the control.

**Figure 5.**
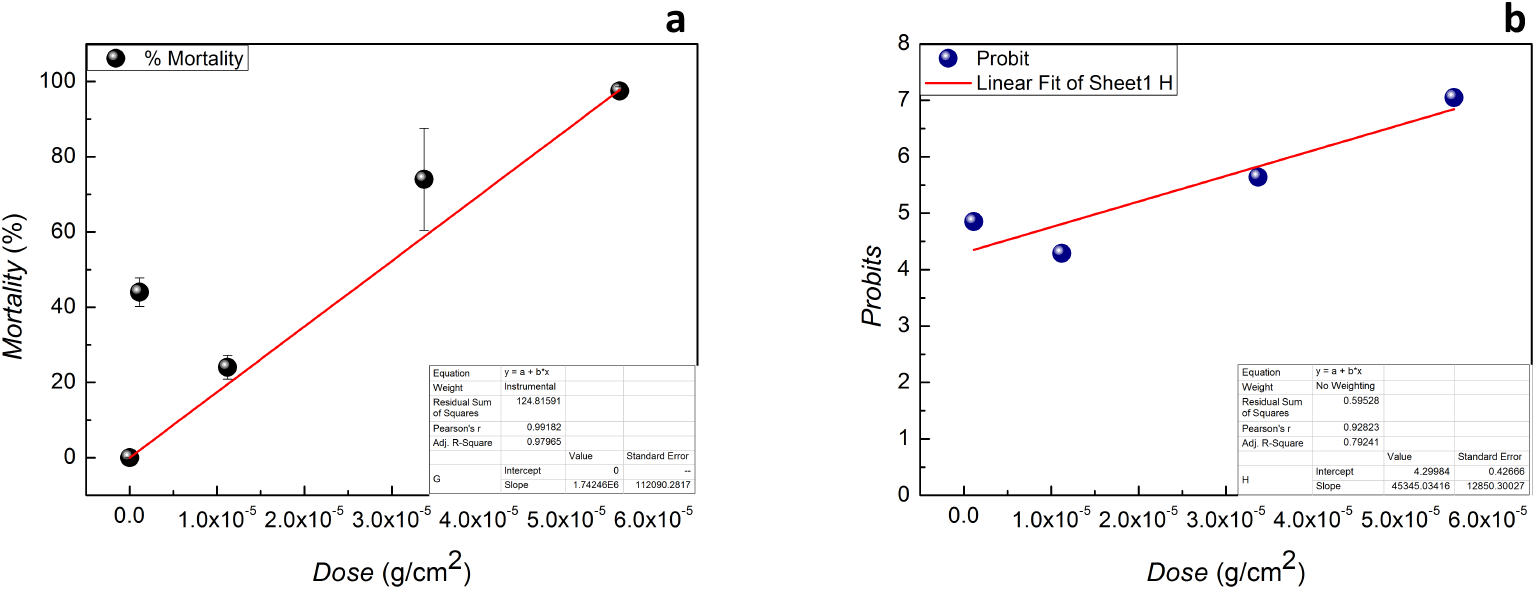
(a) Insect mortality graph, (b) Probit line responses. % values represent mortality levels that are transformed to probit for statistical analysis.

**Table 3.** Dosage level of films, Number of dead insects and total test insects per dose

| Mortality Rate (Cumulative) |  |  |  |  |  |  |  |  |  |  |
| --- | --- | --- | --- | --- | --- | --- | --- | --- | --- | --- |
| Day | Dose (%) |  |  |  |  |  |  |  | Control |  |
|  | 0.1 |  | 1.0 |  | 3.0 |  | 5.0 |  |  |  |
| 1 | 0 | 0 | 0 | 0 | 0 | 0 | 0 | 0 | 0 | 0 |
| 2 | 1 | 1 | 1 | 2 | 4 | 1 | 4 | 5 | 0 | 0 |
| 3 | 1 | 1 | 2 | 3 | 6 | 5 | 9 | 5 | 1 | 0 |
| 4 | 3 | 1 | 2 | 3 | 7 | 9 | 12 | 7 | 1 | 0 |
| 5 | 5 | 1 | 2 | 4 | 8 | 13 | 15 | 7 | 1 | 0 |
| 6 | 6 | 2 | 4 | 4 | 12 | 15 | 20 | 11 | 1 | 0 |
| 7 | 11 | 9 | 6 | 4 | 12 | 17 | 20 | 20 | 1 | 0 |
| 8 | 13 | 10 | 8 | 5 | 20 | 20 | - | - | 4 | 3 |
| 9 | 14 | 12 | 11 | 10 | - | - | - | - | 5 | 3 |
| 10 | 16 | 14 | 14 | 13 | - | - | - | - | 7 | 6 |
| Overall |  |  |  |  |  |  |  |  |  |  |
| Dose (%) | 0.1 |  | 1.0 |  | 3.0 |  | 5.0 |  | Control |  |
| # Mortality per dose <sup>1</sup> | 9.5 |  | 4.5 |  | 14.0 |  | 19.5 |  | 0.5 |  |
| Total # of insects | 20 |  | 20 |  | 20 |  | 20 |  | 20 |  |

<sup>1</sup> # of mortality per dose indicated that average of the three measurements using Abbott's correction; this value was counted on the experiment 7th days.

In addition to *LD50* determination, dietary and jump behavior of grasshopper were also observed throughout the test cycle. Behavioral features (dietary and jump) of the insect population are affected by the DM-HNT loaded LLDPE films. According to restricted diet feeding observations, it was seen that, the feeding amounts decreased after 2 days in 3% and 5% dose sets and after 4 days in 0.1% and 1% dose sets. In addition, insect sets jumping rate decreased after 7 days in control sets, after 6 days in 0.1% dose sets 5 days in 1% dose sets, after 2 days in 3% and 5% sets.

### 3.7. Insecticidal Activity

Insecticidal activity tests of deltamethrin loaded LLDPE films were performed in a greenhouse. Figure 6 (a, b) shows the of DMHNT loaded film application on aphid contaminated trefoils (Fig 6-a, b; emphasized with arrow), and the greenhouse ground that the planter was placed before and after film application (Fig 6-c, d). Comparing the images of foils of contaminated trefoil before and after 10%-DMHNT application, it is observed that insecticide-loaded nanocomposite repel mature aphids (Fig 6-a, b). Moreover, it kills the young aphids (nymph), and thrips (Fig 6-c, d). This implies that the chemical insecticide is still active after loading into HNTs and incorporated into LLDPE films.

**Figure 6.**
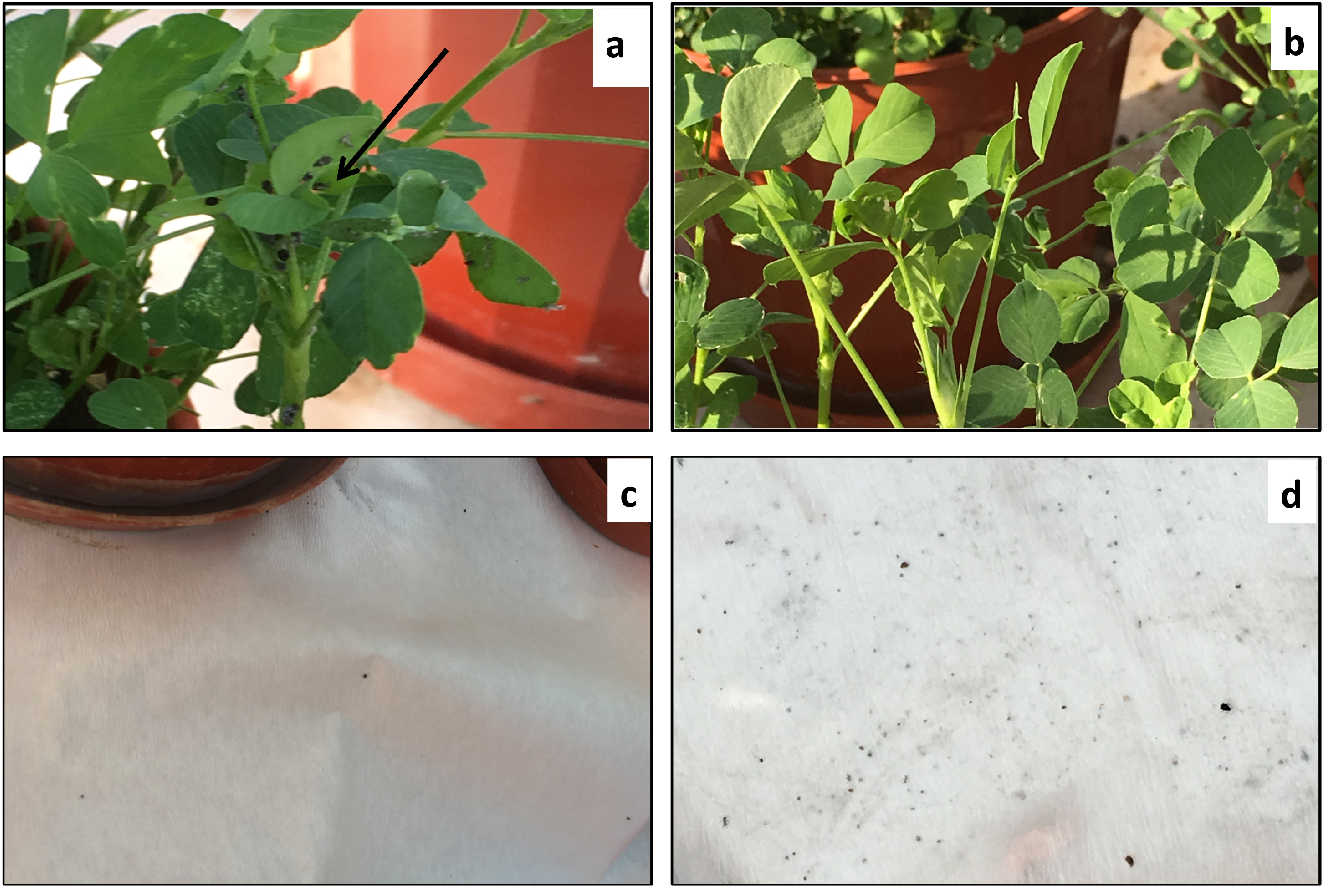
Greenhouse application of 10%-DMHNT film. Foils of aphid contaminated trefoil before (a), and after two days following the 10%-DMHNT treatment (b). Planter ground before (c) and after (d) the treatment under greenhouse conditions.

## 4. Discussion

This study describes the preparation of insecticide-releasing plastic films (LLDPE) that is efficient, cheap, easy to prepare and, suitable to use in agricultural applications. Thermal and morphological traits reveal that DMHNTs were successfully incorporated into LLDPE films. In addition, mechanical properties of LLDPEs are not deteriorated following DMHNT loading, even in the highest concentration. DMHNT incorporation into LLDPE caused an increase in elastic modulus of the films, with a drop in tensile strength, ductility and toughness due to the agglomeration of nano-sized HNTs, which is confirmed by SEM as well. Bioactivities of the prepared films were tested against grasshoppers (*Locusta Migratoria*). Bioassays of the prepared films confirm that the lethal dose for DMHNT-loaded LLDPE films are 1.85*x*10^−5^ g/cm^2^. Insecticidal activities of nanocomposite films were tested in Sabanci University greenhouse, and proven to be effective against two types of pests examined. In this experiment, films were placed around 1 m^2^ of plant’s periphery. Since this test was only applied in close proximity, we can conclude that the resulting nanocomposite film is a good candidate for low tunnel-type greenhouse applications and has a potential application in high tunnels.

## 5. Acknowledgments

The authors thank to Esan Eczacıbaşı A.Ş. for kindly providing untreated halloysite nanotubes, Sabancı University Integrated Manufacturing Technologies Research and Application Center (SU-IMC) for providing facility resources for composite film preparation, and Mustafa Atilla Yazici for their invaluable help in greenhouse tests.

## 6. Patents

“Kontrollü Salınım Yapan Pestisit Yüklü Sera Ortüsü” Turkish Patent and Trademark Office Application No: 10-18774-A

